# Developmental rate displays effects of inheritance but not of sex in inter-population hybrids of *Tigriopus californicus*

**DOI:** 10.1101/2022.09.13.507602

**Authors:** Timothy M. Healy, Alexis Cody Hargadon, Ronald S. Burton

## Abstract

Coevolved interactions between mitochondrial-encoded and nuclear-encoded genes within populations can be disrupted by inter-population hybridization resulting in reduced hybrid fitness. This hybrid breakdown may be an important factor contributing to reproductive isolation between populations or species, and strong selection among hybrids to maintain compatible mitonuclear genotypes occurs in at least some species. Despite potentially differential consequences of mitonuclear incompatibilities in females and males due to maternal inheritance of the mitochondrial genome, the extent to which phenotypic variation associated with hybrid breakdown is sex-specific and heritable remains unresolved. Here we present two experiments investigating variation in developmental rate among reciprocal inter-population hybrids of the intertidal copepod *Tigriopus californicus*. Developmental rate is a proxy for fitness in this species that is substantially influenced by variation in mitonuclear compatibility among hybrids. First, we show that F_2_ hybrid developmental rate is the same in females and males, suggesting that effects of mitonuclear incompatibilities on this trait are likely experienced equally by the two sexes. Second, we demonstrate that variation in developmental rate among F_3_ hybrids is heritable; times to copepodid metamorphosis of F_4_ offspring of fast-developing F_3_ parents (12.25 ± 0.05 d, μ ± SEM) were significantly faster than those of F_4_ offspring of slow-developing parents (14.58 ± 0.05 d). Taken together, these results provide evidence for strong effects of mitonuclear interactions across generations of hybrid eukaryotes with no differences between the sexes, and support key roles of mitonuclear incompatibility in hybrid breakdown and reproductive isolation.

## Introduction

In eukaryotic organisms, the functions of mitochondrial-encoded proteins and RNAs (13 proteins, 2 rRNAs and 22 tRNAs in bilaterian animals; Levin et al., 2014) rely on essential interactions with the products of nuclear-encoded genes (Rand et al., 2004; Lane, 2005; Burton & Barreto, 2012; Hill, 2015). These mitonuclear interactions are key for mitochondrial performance, and consequently incompatibilities between the two genomes have the potential to result in loss of fitness (Burton et al., 2006, 2013; Hill et al., 2019). Compatibility within a population or species is typically maintained by inter-genomic coevolution (Rand et al., 2004; Hill, 2015; Burton et al., 2013), but because mitochondrial DNA evolves at relatively high rates (Lynch, 1997; Wallace, 2010), coevolved gene interactions may often be species- or population-specific. Therefore, compatible mitonuclear interactions can be disrupted due to the reassortment of nuclear alleles and mitochondrial genotypes in hybrid organisms resulting in hybrid breakdown and potentially contributing to reproductive isolation between taxa (Burton & Barreto, 2012; Hill et al., 2019). However, for mitonuclear interactions to be major factors underlying early-stage reproductive isolation, and potentially speciation (Gershoni et al., 2009; Sloan et al., 2017; Hill, 2017; Burton, 2022), incompatibilities need to result in negative fitness effects that are sufficient to create strong selection pressures favoring compatible mitonuclear genotypes (Sloan et al., 2017; Hill et al., 2019).

Substantial effects of inter-genomic interactions on hybrid fitness have been observed in several species (e.g., Ellison & Burton, 2008b; Meiklejohn et al., 2013; Moran et al., 2022), and consistent with strong selection for mitonuclear compatibility, key roles of incompatibilities underlying variation in fitness-related traits among F_2_ hybrids have recently been identified (Healy & Burton, 2020; Han & Barreto, 2021). The heritability of this trait variation among hybrid individuals is potentially another important factor determining if mitonuclear incompatibilities can create strong barriers for gene flow between populations, and to our knowledge, the transmission of incompatibilities across generations has not been directly assessed. Additionally, it is possible for the effects of mitonuclear incompatibilities to vary between females and males (Jelić et al., 2015, Mossman et al., 2016b; Đorđević et al., 2017; Hoekstra et al., 2018, Carnegie et al., 2021; Erić et al., 2022), which may be the result of genetic interactions involving sex determining loci (e.g., heterologous sex chromosomes; Lopez et al., 2021), or of the predominantly maternal inheritance of mitochondrial DNA in metazoans (e.g., Giles et al., 1980). For example, the ‘mother’s curse’ hypothesis posits that mutations in mitochondrial DNA causing beneficial interactions in females will accumulate, even if they cause negative interactions in males, due to the lack of paternal transmission of mitochondrial DNA (Frank & Hurst, 1996; Gemmell et al., 2004). Consistent with these expectations, effects of mitonuclear interactions aligning with the ‘mother’s curse’ have been observed for many traits (Rand et al., 2001; Camus et al., 2012; Milot et al., 2017; Carnegie et al., 2021, although see Mossman et al., 2016a, 2016b, 2017; Eyre-Walker, 2017; Rand & Mossman, 2020). Taken together, it is clear not only that mitonuclear incompatibilities can have substantial fitness effects, but also that resolving the impacts of heritability and sex on traits associated with these interactions is critical for understanding the role of mitonuclear incompatibility in outbreeding depression, early-stage reproductive isolation and speciation.

*Tigriopus californicus* is a species of intertidal copepod that is ideal for the study of mitonuclear interactions. These copepods inhabit supralittoral tidepools found on rocky outcrops along the Pacific coast of North America from Baja California, Mexico to Alaska, USA. There is virtually no migration of *T. californicus* between rocky outcrops (Burton & Feldman, 1981; Burton, 1997), resulting in extremely high genetic divergence among populations of this species, particularly in the mitochondrial genome (Burton et al., 2007; Barreto et al., 2018). However, laboratory crosses between populations generate offspring that are both viable and fertile (e.g., Burton, 1986), and previously published results in *T. californicus* have demonstrated hybrid breakdown from the F_2_ generation onwards that was consistent with the effects of mitonuclear incompatibilities on fitness-related traits, such as fecundity, viability, hatching success, metamorphosis success, developmental rate and mitochondrial performance (Burton, 1986, 1987, 1990; Edmands, 1999; Edmands & Burton, 1999; Willett & Burton, 2001, 2003; Ellison & Burton, 2006, 2008a, 2008b; Healy & Burton, 2020; Han & Barreto, 2021). Several studies have investigated the genetic mechanisms associated with these phenotypic effects, and signatures of both nuclear-nuclear and mitonuclear incompatibilities underlying hybrid breakdown have been detected, although mitonuclear effects tend to dominate overall (Edmands et al., 2009; Pritchard et al., 2011; Foley et al., 2013; Lima et al., 2019; Healy & Burton, 2020; Han & Barreto, 2021; Pereira et al., 2021). This dominant role of mitonuclear interactions is particularly evident for variation in developmental rate among F_2_ hybrids, and there is strong selection for compatible mitonuclear genotypes in fast-developing hybrids (Healy & Burton, 2020; Han & Barreto, 2021). Previous estimates for the times necessary to reach adulthood in inter-population *T. californicus* hybrids suggest developmental rate is equivalent in both sexes (Burton, 1990), which is unlike many other traits in this species (Willett & Burton, 2001; Foley et al., 2013; Flanagan et al., 2021; Li et al., 2022; Watson et al., 2022). However, the potential effects of sex throughout development are not well characterized, and the extent to which mitonuclear effects on developmental rate are inherited across hybrid generations in unknown.

To address these knowledge gaps, we present the results of two experiments that (1) investigate the effects of sex on developmental rate in *T. californicus* inter-population hybrids, and (2) assess the inheritance of variation in developmental rate between hybrid generations. In particular, we focus on these questions: Does developmental rate vary between female and male F_2_ hybrids throughout development? Is variation in developmental rate among hybrids dependent on mitochondrial genotype? Is fast or slow development among F_3_ hybrids inherited by their F_4_ offspring? Among F_4_ hybrids, are there associations between developmental rate and mitochondrial performance?

## Materials & methods

### Copepod collection and laboratory acclimation

*T. californicus* were collected from splashpools in the intertidal zone in San Diego, California (SD; 32° 45’ N, 117° 15’ W), and Santa Cruz, California (SC; 36° 56’ N, 122° 02’ W) in the summer of 2018. Copepods were transferred into 1 L plastic bottles containing water from their tidepools using large plastic pipettes, and transported to Scripps Institution of Oceanography, University of California San Diego within 24 h. Collected copepods were split among several 250 mL laboratory cultures in 400 mL glass beakers that were held in incubators set at 20 °C and 12h: 12h light:dark, and cultures were maintained with filtered (0.44 μm) natural seawater (35 psu). Powdered spirulina (Salt Creek, Inc., South Salt Lake City, UT, USA) and ground TetraMin^®^ Tropical Flakes (Spectrum Brands Pet LLC, Blacksburg, VA, USA) were added as food to the cultures once per week, and copepods also consumed natural algal growth within their cultures. Laboratory cultures were maintained for a minimum of three months (i.e., ~3 generations) prior to the start of experiments.

### Developmental rate in female and male F_2_ hybrids

Virgin SD and SC females were obtained by splitting mate-guarding pairs with a fine needle (Burton et al., 1981; Burton, 1985), and reciprocal crosses between the populations, SD♀ x SC♂ (SDxSC) or SC♀ x SD♂ (SCxSD), were initiated by placing 40 virgin females from one population in ~200 mL of filtered seawater in a 2.5 × 15 cm petri dish with 40 males from the other population. Females and males paired haphazardly, and when gravid P_0_ females (i.e., females carrying an egg sac) were observed they were moved to a new dish. All dishes were maintained under the same conditions as the laboratory cultures. Egg sacs hatched naturally in the new dishes, and when F_1_ hybrid offspring were visible to the naked eye, the P_0_ females were removed to avoid overlapping generations in the dishes. After maturation, F_1_ adults paired and mated haphazardly, and F_2_ developmental rate trials were initiated one week after gravid F_1_ females were observed. This avoided using only the fastest-developing F_1_ hybrids as parents in the F_2_ trials while still preventing any F_2_ hybrids from reaching adulthood in the F_1_ dishes.

Mature (i.e., red) egg sacs were removed from F_1_ gravid females using a fine needle and placed individually in wells of Falcon^®^ 6-well plates (Thermo Fisher Scientific, Waltham, MA, USA). This was done for 10 SDxSC and 10 SCxSD egg sacs. Nauplii (i.e., larval copepods) hatched from the egg sacs overnight, and development was tracked daily by observation through a stereo microscope as in Healy & Burton (2020). *Tigriopus sp.* development involves a distinct metamorphosis from the last naupliar stage (N6) to the first copepodid stage (C1; Raisuddin et al., 2007), and developmental rate can be assessed by the time from hatch to metamorphosis. As copepodids appeared, they were transferred individually into wells of Falcon^®^ 24-well plates (Thermo Fisher Scientific), and developmental progress through the copepodid stages (C1, C2, C3, C4, C5 and adult) was monitored as described by Tsuboko-Ishii & Burton (2018). The times between stage transitions were recorded for each individual, and developmental rate was scored for a total of 478 F_2_ hybrids (273 SDxSC and 205 SCxSD). Throughout these developmental trials the nauplii and copepodids were fed every other day by the addition of powdered spirulina to the 6- or 24-well plates.

### Developmental rate in F_3_ hybrids and their F_4_ offspring

In a separate experiment, three hybrid lines for both reciprocal crosses between the SD and SC populations (i.e., six lines total) were initiated as described above for the F_2_ experiment. However, the initial crosses between the SD and SC populations involved 50 virgin females and 50 males for each line, and rather than dissecting F_2_ egg sacs from gravid F_1_ females to score developmental rate, 144 mature F_2_ egg sacs were deposited into new 2.5 × 15 cm petri dishes (one per line) and allowed to hatch overnight. Within these dishes, the F_2_ hybrids matured, haphazardly paired and mated. Gravid F_2_ females were transferred to a new dish for each line, and F_3_ developmental rate trials began approximately two weeks after the transfer of the first gravid F_2_ female to the new dish. The increase in interval between the F_2_ gravid female transfers and the collection of egg sacs compared to the interval for F_1_ females (see above) was to account for generally high levels of variation in developmental rate among F_2_ hybrids (e.g., Healy & Burton, 2020).

Mature F_3_ egg sacs were dissected from 60 gravid F_2_ females for each line, and were placed individually into wells of Falcon^®^ 6-well plates to allow the days post hatch (dph) to C1 metamorphosis to be scored for each F_3_ offspring. In total, developmental rate was determined for 11,263 F_3_ copepodids with between 1,360 and 2,537 individuals scored per line. The F_3_ copepodids were divided into two developmental rate groups for each line: fast developers that metamorphosed ≤ 10 dph and slow developers that metamorphosed ≥ 17 dph. Each group was established in its own 2.5 × 15 cm petri dish, and the number of copepodids establishing each group ranged from 77 to 597. As for the F_2_ hybrids, the F_3_ offspring matured and mated haphazardly, and gravid F_3_ females were transferred to a new dish when they were observed. Two weeks after the initial transfers of the gravid F_3_ females, 30 mature F_4_ egg sacs were collected for each group (i.e., 12 groups for each line x F_3_ parental developmental rate combination) from the gravid females, and time to C1 metamorphosis was scored for the individual F_4_ offspring. Developmental rate was assessed for 672 to 1157 F_4_ copepodids per group with 10,018 F_4_ individuals scored overall.

### Mitochondrial ATP synthesis rates in F_4_ hybrids

ATP synthesis rates were measured *in vitro* for mitochondria isolated from the F_4_ *T. californicus* hybrids using an approach similar to those presented in Harada et al. (2019), Healy et al. (2019) and Healy and Burton (2020). In brief, mitochondria were isolated from pools of 15 F_4_ copepods from each line x F_3_ parental developmental rate group once the copepods had reached adulthood. Copepods were homogenized by hand in 800 μL of ice-cold buffer (400 mmol L^-1^ sucrose, 100 mmol L^-1^ KCl, 70 mmol L^-1^ HEPES, 6 mmol L^-1^ EGTA, 3 mmol L^-1^ EDTA, 1% w/v BSA, pH 7.6) with a Teflon-on-glass homogenizer, and mitochondria were isolated by differential centrifugation, resuspended in 130 μL of an ice-cold assay buffer (560 mmol L^-1^ sucrose, 100 mmol L^-1^ KCl, 70 mmol L^-1^ HEPES, 10 mmol L^-1^ KH_2_PO_4_, pH 7.6), supplied with saturating concentrations of ETS complex I or II substrates and allowed to synthesize ATP under saturating conditions for 10 min at 20 °C. Initial and final concentrations of ATP for each reaction were determined with CellTiter-Glo (Promega, Madison, WI, USA), and reactions were performed with three sets of substrates for each sample. Two of the sets of substrates led to donation of electrons to complex I: (1) 5 mmol L^-1^ pyruvate and 2 mmol L^-1^ malate (PM) and (2) 10 mmol L^-1^ glutamate and 2 mmol L^-1^ malate (GM), and one set led to donation of electrons to complex II: 10 mmol L^-1^ succinate (S). All sets of substrates also included 1 mmol L^-1^ ADP. In general, ATP synthesis reactions were performed for 6 pools of adult females and 6 pools of adult males from each group of F_4_ hybrids. However, many F_4_ individuals from these groups were allocated to other experiments in our laboratory, and some groups demonstrated strongly skewed sex ratios (e.g., Alexander et al., 2015). Consequently, for the line A males from fast-developing F_3_ parents and for the line C males from slow-developing F_3_ parents only 5 pools were available for ATP synthesis reactions, and for males from line F with slow-developing F_3_ parents ATP synthesis reactions were only possible for 36 individuals. In this last case, the males were homogenized in 3 pools of 12, resuspension volumes were adjusted proportionally, and GM-substrate reactions were not conducted. Thus, the PM and S ATP synthesis rates for this group (line F males from slow-developing F_3_ parents) were stoichiometrically equivalent to the PM and S measurements for the other pools of copepods; however, given the protocol adjustments that were necessary and the relatively small sample size for this group (n = 3), comparisons between this group and others should be interpreted with some caution.

ATP synthesis rates were normalized to citrate synthase (CS) reaction rates, which were measured *in vitro* for each mitochondrial isolation. CS is a nuclear-encoded enzyme that functions within the tricarboxylic acid cycle, and is commonly used as an index of mitochondrial amount to normalize oxidative phosphorylation reaction rates (e.g., Gnaiger, 2020). CS assays were run in duplicate using 5 μL of the mitochondrial isolations in each replicate. Reactions were conducted in flat-bottomed 96-well plates (Corning, Glendale, AZ, USA) at a final volume of 120 μL with reactant and buffer component concentrations as follows: 50 mmol L^-1^ Tris pH 8.0, 0.1 mmol L^-1^ 5,5’-Dithiobis(2-nitrobenzoic acid), 0.3 mmol L^-1^ Acetyl CoA, 0.5 mmol L^-1^ oxaloacetate and 0.1% vol/vol Triton X-100 (Spinazzi et al., 2012). Prior to the addition of oxaloacetate, background reaction rates were measured by tracking the change in absorbance at 412 nm over 5 min using a SpectraMax^®^ iD3 Multi-Mode Microplate Reader (Molecular Devices, LLC., San Jose, CA, USA). After the addition of oxaloacetate to start the reaction, the change in absorbance at 412 nm was again measured for 5 min, and reaction rates were determined by regressing the change in absorbance against time (in general R^2^ ≥ 0.99). Final CS reaction rates were calculated by subtracting the background rate from the reaction rate (i.e., with oxaloacetate) for each replicate, and then taking the average value of the duplicates for each mitochondrial isolation. ATP synthesis rates were normalized to the CS reaction rates by division.

### Statistical analyses

All statistical tests were performed in *R* v4.2.0 (R Core Team, 2022) with α = 0.05. The majority of analyses in the current study were performed using linear mixed-effects models implemented with the *lmerTest* package v3.1.3 (Kuznetsova et al., 2017). For effects of sex on F_2_ development, cross and sex were fixed factors, and egg sac was a random factor. For effects of cross on F_3_ developmental rate, cross was a fixed factor, and there were nested random factors with egg sac nested within line. For effects of F_3_ parental developmental rate on F_4_ developmental rate, an overall model had F_3_ developmental rate and cross as fixed factors, and egg sac nested within group (i.e., line x F_3_ developmental rate groups) as random factors. Following the overall model, line-specific models were fit with F_3_ parental developmental rate as a fixed factor and egg sac as a random factor. For variation in developmental rate between F_3_ and F_4_ hybrids, cross and line were fixed factors, and egg sac was nested within line as random factors. For F_4_ ATP synthesis rates, cross, sex and F_3_ parental developmental rate were fixed factors and line was a random factor. In general, final models were simplified by removing non-significant interaction terms hierarchically by the order of interactions, and developmental rate data were log transformed prior to model fitting. Realized heritabilities (h^2^) for developmental rate were calculated for each line with the breeder’s equation: R = h^2^ x S (Lush, 1937). In the current study, S was equal to the average developmental rate for the fast- or slow-developing F_3_ parents minus the average F_3_ developmental rate, and R was equal to the average F_4_ developmental rate minus the average F_3_ developmental rate. Relationships between the hatch to C1, and C1 to adult development rates were assessed with a linear model for variation in C1 to adult developmental rate that had hatch to C1 developmental rate as a continuous factor and cross as a fixed factor. Potential effects of density dependence on developmental rate were assessed in F_3_ and F_4_ hybrids using linear models for variation in the average developmental rate for an egg sac that had the number of copepodids from an egg sac as a continuous factor and either line or line x F_3_ parental developmental rate group as a fixed factor (F_3_ and F_4_ models, respectively). For the F_4_ hybrids, additional models were fit within each line x F_3_ parental developmental rate group with the number of copepodids from an egg sac as a continuous factor followed by analysis of variance tests utilizing the *R* package *car* v3.0-13 (Fox & Weisberg, 2019).

## Results

### Developmental rates in female and male F_2_ hybrids

There was no variation in developmental rate between female and male F_2_ hybrids when assessed by time from hatch to copepodid stage 1 (*p* = 0.51; Figure 1a), copepodid stage 1 to adult (*p* = 0.20; Figure 1b), or hatch to adult (*p* = 0.62; Figure 1c), and variance in developmental rate did not differ between the sexes for any measure of developmental rate in either cross (i.e., in SDxSC or SCxSD hybrids; Bartlett test *p* ≥ 0.07). There were also no effects of cross on any of these metrics (*p* ≥ 0.44). Interestingly, there was no significant relationship between the times from hatch to C1 and from C1 to adult across egg sacs (*p* = 0.15; Figure 2). In general, approximately 2 to 2.5 d were spent at each copepodid stage with the time at C5 tending to be slightly longer than times spent at other stages (Table 1), and similar to overall developmental rate, there were no effects of sex on rate of development at any specific copepodid stage (*p* ≥ 0.12). At most stages, there were no differences in developmental rate between SDxSC and SCxSD copepodids (*p* ≥ 0.66), but SDxSC copepodids spent less time at stage 1 than SCxSD copepodids did (*p* = 0.009).

**Figure 1.**
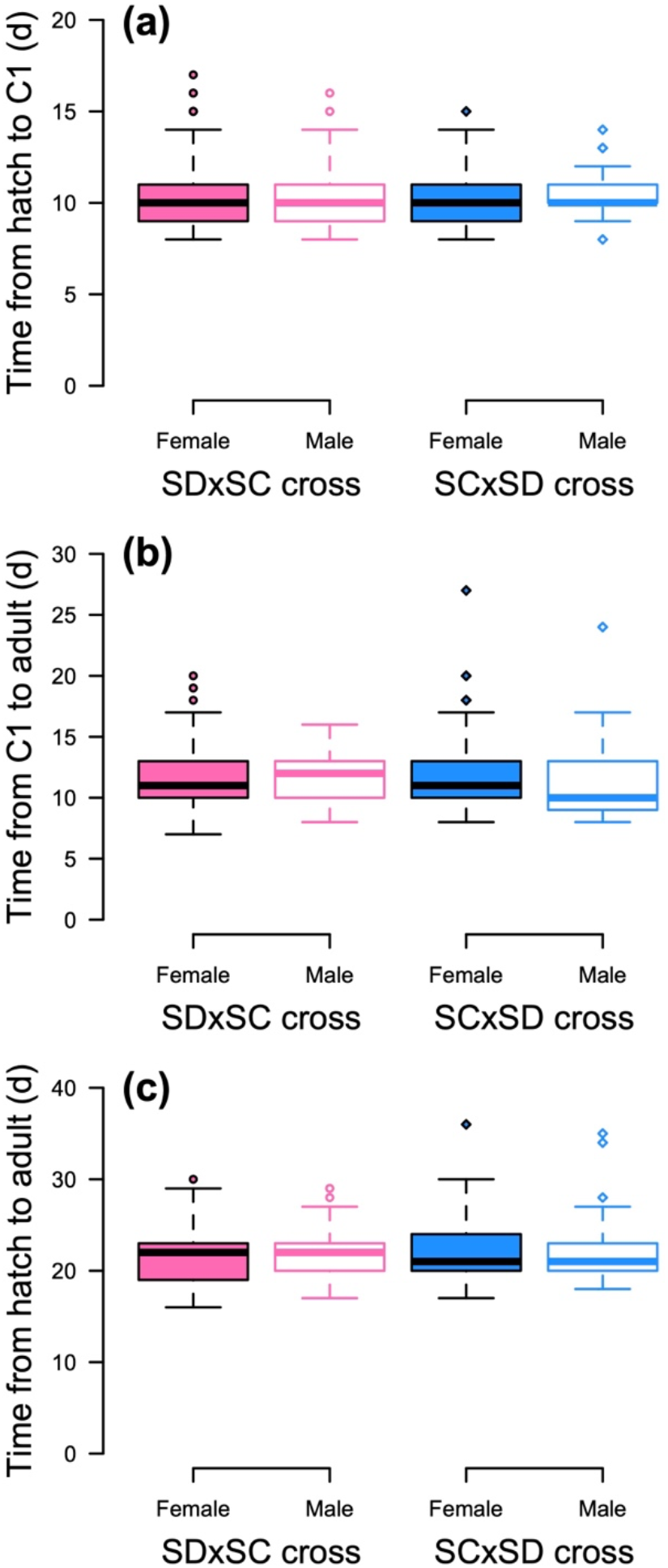
Box plots of developmental rate from (a) hatch to stage 1 copepodid (C1), (b) C1 to adult, and (c) hatch to adult for female (filled boxes) and male (open boxes) F_2_ *T. californicus* reciprocal hybrids (SDxSC – pink; SCxSD – blue). No significant effects of sex or cross were detected for any of the measurements of developmental rate.

**Figure 2.**
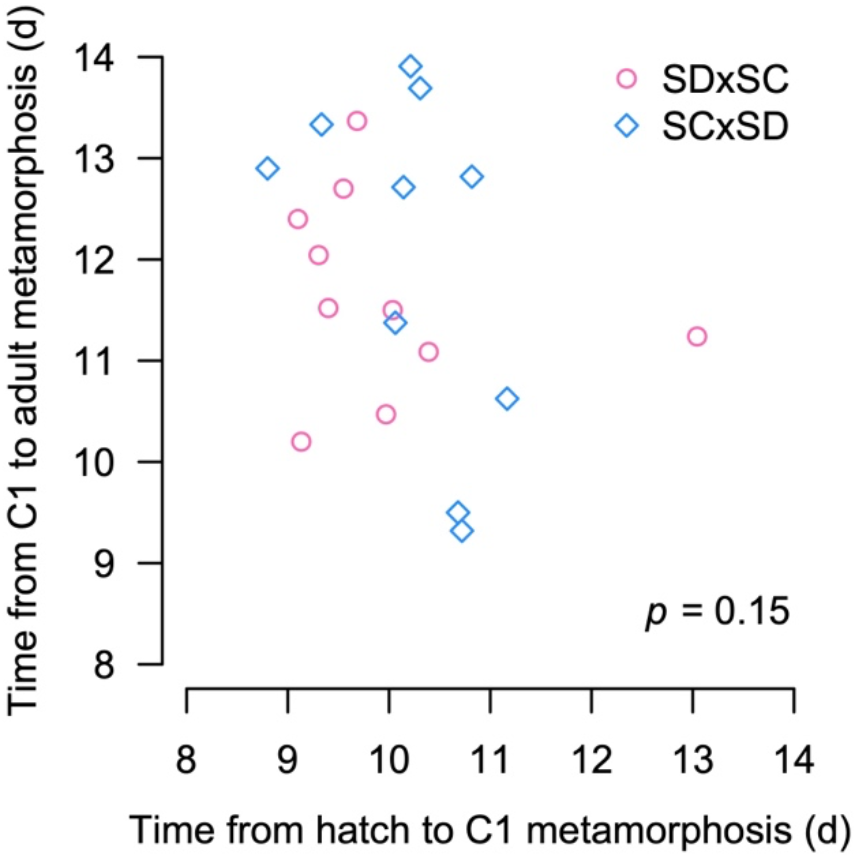
Average egg sac developmental rate from copepodid stage 1 (C1) to adult (y-axis) plotted against developmental rate from hatch to C1 (x-axis). Data are presented as time to adult or C1 metamorphosis, respectively (SDxSC egg sacs: pink, circles; SCxSD egg sacs: blue, diamonds). The *p*-value for an effect of hatch to C1 developmental rate on C1 to adult developmental rate is displayed on the graph.

**Table 1.** Time to metamorphosis at each copepodid stage during *T. californicus* development in female and male F_2_ inter-population SDxSC and SCxSD hybrids.

| Developmental<br>stage<br>transition | Time to metamorphosis (d) ($\mu \pm \text{SEM}$) | | | |
| --- | --- | --- | --- | --- |
|  | SDxSC cross |  | SCxSD cross |  |
|  | Female | Male | Female | Male |
| *C1 to C2*<sup>†</sup> | 1.85 ± 0.05 | 1.80 ± 0.07 | 2.23 ± 0.07 | 2.11 ± 0.11 |
| *C2 to C3* | 2.39 ± 0.07 | 2.26 ± 0.09 | 2.21 ± 0.06 | 2.22 ± 0.09 |
| *C3 to C4* | 2.40 ± 0.08 | 2.41 ± 0.08 | 2.25 ± 0.06 | 2.38 ± 0.19 |
| *C4 to C5* | 2.26 ± 0.07 | 2.28 ± 0.09 | 2.14 ± 0.08 | 2.07 ± 0.14 |
| *C5 to A* | 2.69 ± 0.06 | 2.95 ± 0.11 | 2.61 ± 0.12 | 2.65 ± 0.16 |

<sup>†</sup> significant difference between SDxSC and SCxSD ( $p = 0.009$ )

### Inheritance of developmental rate between F_3_ and F_4_ hybrids

Across the F_3_ lines in our study metamorphosis to the C1 stage was observed from 7 to 45 dph, and there was no significant difference in developmental rate between the SDxSC and SCxSD lines (*p* = 0.09) with only a small trend for faster development in SDxSC (12.14 ± 0.05 d, μ ± SEM) than in SCxSD (13.49 ± 0.04 d; Figure 3). The F_3_ copepodids were grouped into fast and slow developers (≤ 10 and ≥ 17 dph, respectively), and among lines, 9-33 egg sacs contributed to only the fast-developing groups, 4-25 contributed to only the slow-developing groups and 18-47 contributed to both groups.

**Figure 3.**
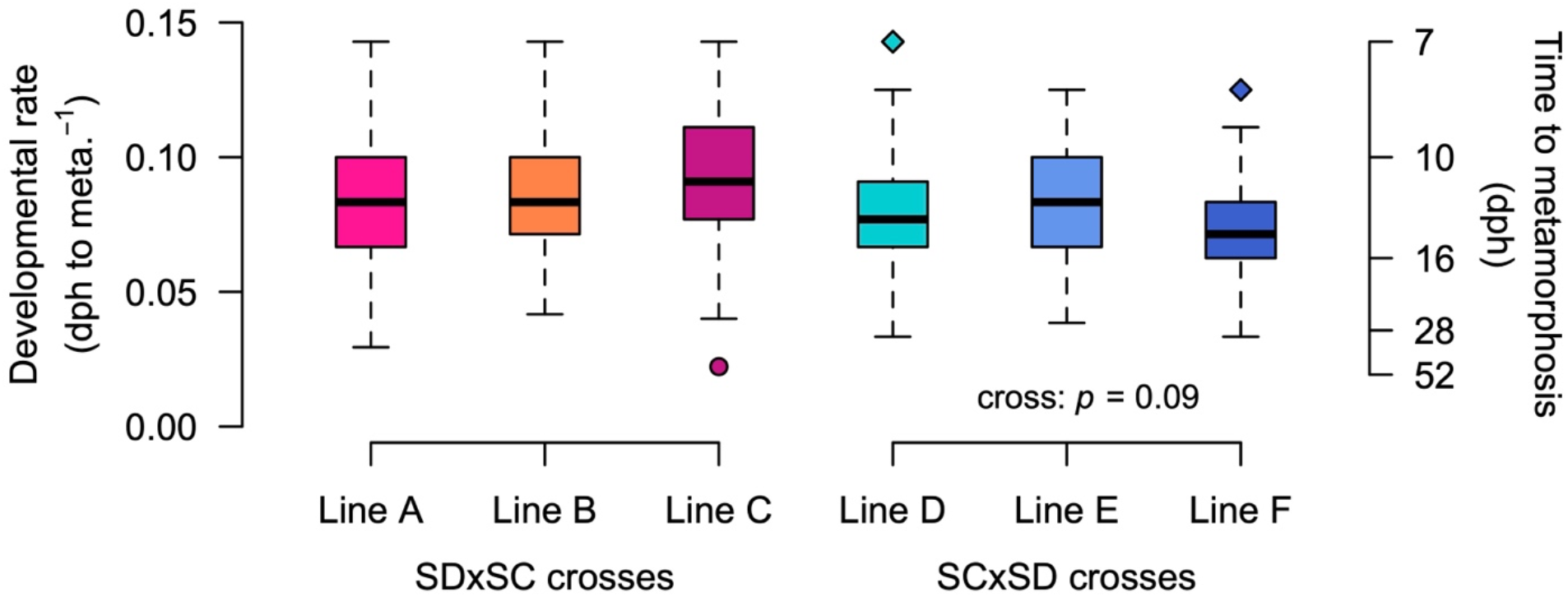
Box plots of developmental rate from hatch to stage 1 copepodid (C1) metamorphosis for F_3_ *T. californicus* reciprocal hybrids (SDxSC, lines A-C – warm colors, circles; SCxSD, lines D-F – cool colors, diamonds). Developmental rates are shown as a rate (left axis) and as a time to metamorphosis (right axis). The *p*-value for potential variation between the SDxSC and SCxSD crosses is shown on the graph.

Developmental rates in the F_4_ offspring of parents from the fast- and slow-developing F_3_ groups were not affected by cross (i.e., SDxSC versus SCxSD; *p* = 0.11), but were affected by the developmental rate group of their F_3_ parents (*p* = 0.005; Figure 4). For all lines, the median developmental rates of the F_4_ offspring of the fast-developing F_3_ hybrids were higher than the median developmental rates of the F_4_ offpsring of the slow-developing F_3_ hybrids, and in all but SCxSD line E, there were significant differences between offspring from the fast- and slow-developing F_3_ groups (*p* = 0.46 for SCxSD line E and *p* ≤ 0.015 for all other lines). Developmental rates tended to be faster in F_4_ offspring from fast-developing F_3_ parents than in the F_3_ generation on average, but these differences were not significant (*p* = 0.12; Figure 5a). In contrast, the developmental rates of F_4_ offspring from slow-developing F_3_ parents were significantly slower than the F_3_ generation on average (*p* = 9.4 × 10^-14^; Figure 5b). Realized heritabilities for developmental rate between the F_3_ and F_4_ generations were 0.16 ± 0.10 and 0.29 ± 0.06 (μ ± SEM) for the fast and slow developers, respectively (Table 2), or 0.24 ± 0.06 and 0.27 ± 0.08 if only the lines displaying significant differences between the offspring of fast or slow developers were considered (lines A, B, C, D and F).

**Figure 4.**
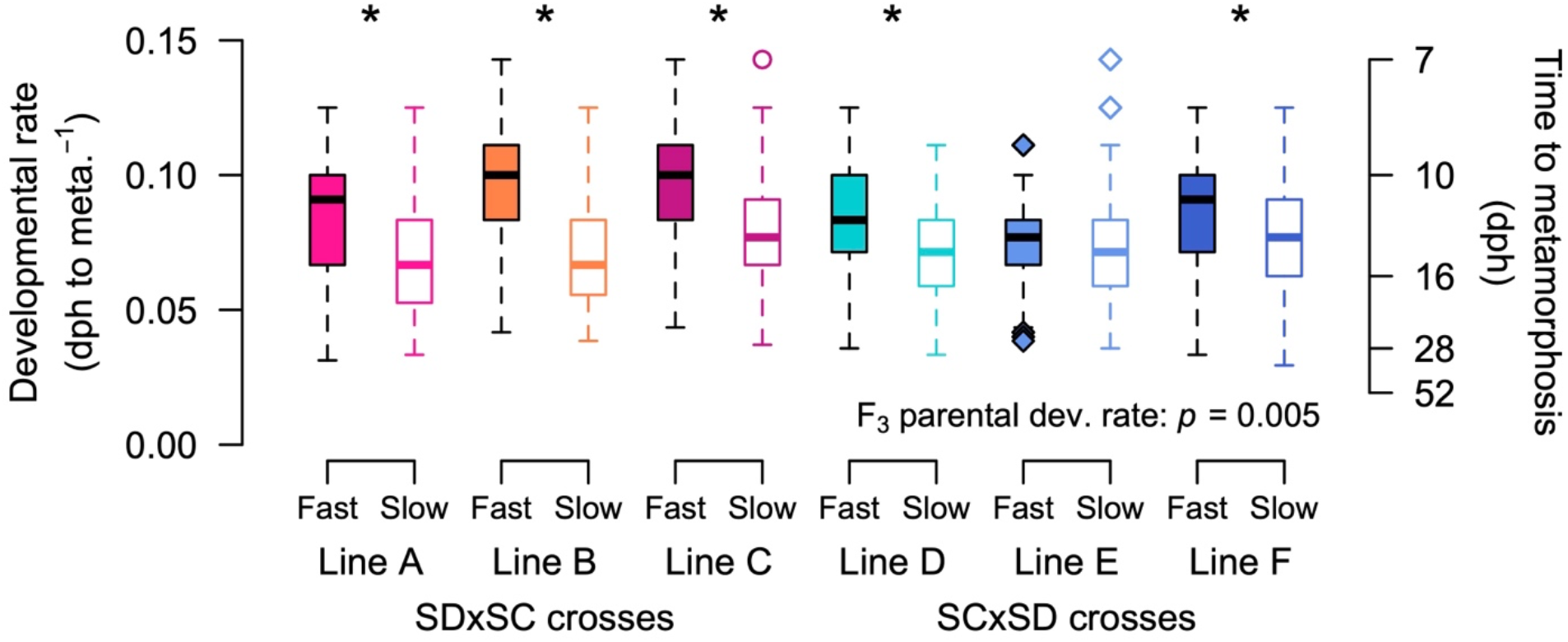
Box plots of developmental rate from hatch to stage 1 copepodid (C1) metamorphosis for F_4_ *T. californicus* reciprocal hybrids (SDxSC, lines A-C – warm colors, circles; SCxSD, lines D-F – cool colors, diamonds) that were offspring of ‘Fast’- or ‘Slow’-developing F_3_ parents (filled or open boxes, respectively). Developmental rates are shown as a rate (left axis) and as a time to metamorphosis (right axis). The *p*-value for an effect of F_3_ parental developmental rate in general is shown on the graph, and asterisks indicate significant effects of F_3_ parental developmental rate that were detected by line-specific tests.

**Figure 5.**
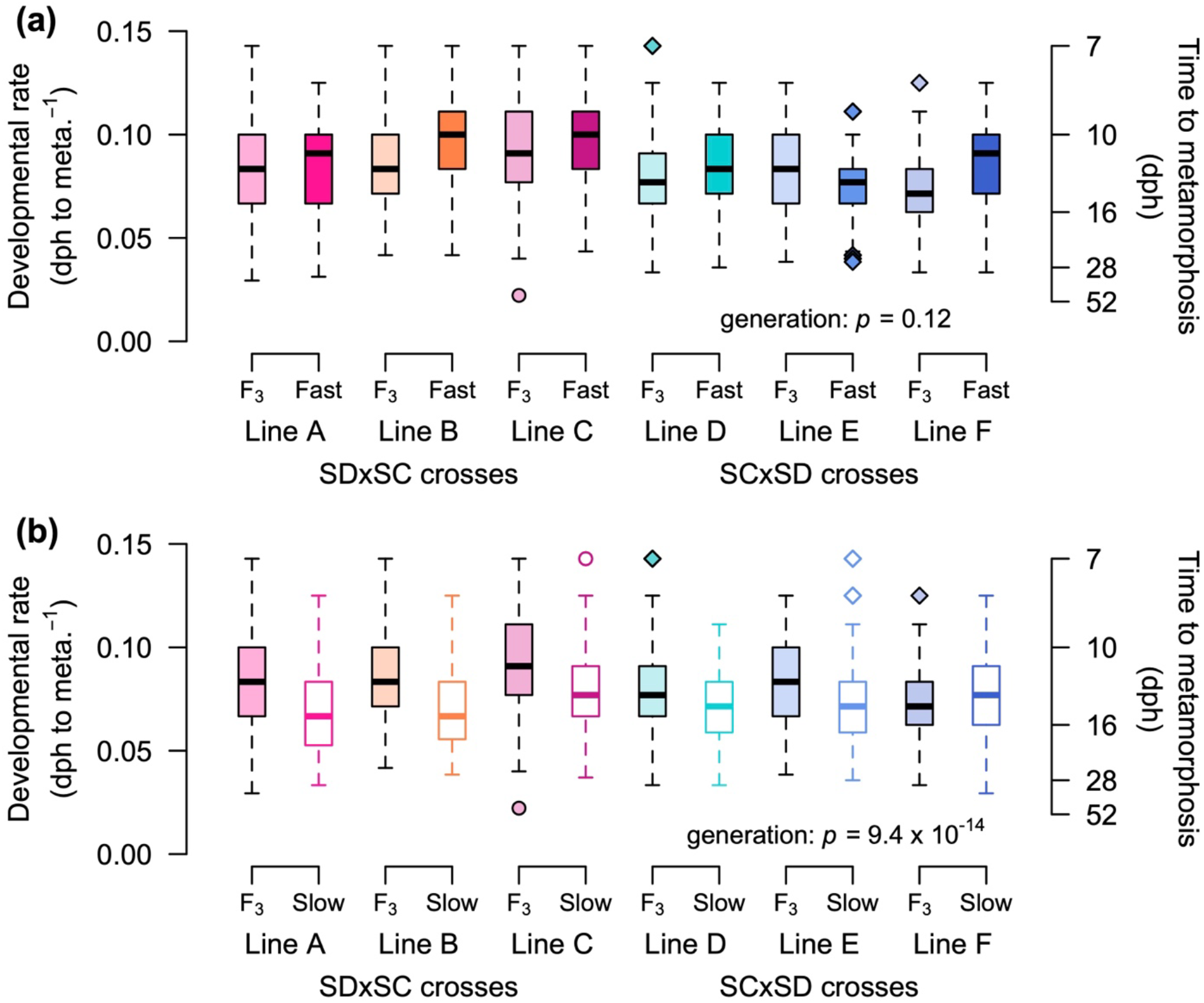
Box plots of developmental rate in F_3_ inter-populations hybrids (‘F_3_’ – filled boxes, faded colors) compared to developmental rate in F_4_ offspring from fast-developing F_3_ parents (a: ‘Fast’ – filled boxes, saturated colors) or from slow-developing F_3_ parents (b: ‘Slow’ – open boxes). Data for SDxSC crosses are shown in warm colors (lines A-C, circles) and SCxSD crosses are shown in cool colors (lines D-F, diamonds). Developmental rates are shown as a rate (left axis) and a time to copepodid stage 1 metamorphosis (right axis). The *p*-values for variation between the F_3_ and F_4_ generations are shown in each panel

**Table 2.** Realized heritabilities (h^2^) for developmental rate between the F_3_ and F_4_ generations for fast- and slow-developing hybrid *T. californicus*.

| Cross | Line | Realized heritability ($h^2$; F<sub>3</sub> to F<sub>4</sub> generation) | |
| --- | --- | --- | --- |
|  |  | Fast developers | Slow developers |
| SDxSC | *A* | 0.06 | 0.39 |
|  | *B* | 0.29 | 0.45 |
|  | *C* | 0.28 | 0.24 |
| SCxSD | *D* | 0.18 | 0.26 |
|  | *E* | -0.26 | 0.37 |
|  | *F* | 0.41 | 0.01 |

In general, egg sacs of females from the slow-developing F_3_ groups produced higher numbers of copepodids than eggs sacs of females from the fast-developing groups did. As a result, we assessed if developmental rates across egg sacs were consistent with effects of density dependence. For the F_3_ copepodids (Figure 6a-f), average developmental rates across egg sacs were not affected by line (*p* = 0.17) or by the number of copepodids that metamorphosed from an egg sac (*p* = 0.28), and there was no interaction between these effects (*p* = 0.58). For the F_4_ copepodids (Figure 6g-l), there was significant effect of line x F_3_ parental developmental rate group (*p* = 3.8 × 10^-4^), and no significant effect of copepodid number per egg sac (*p* = 0.18). However, in this case, there was a significant interaction between these factors (*p* = 1.4 × 10^-3^). Thus, effects of the number of copepodids per egg sac were analyzed separately for each line x F_3_ parental developmental rate group. For 8 of the 12 groups, there was no significant effect of copepodid number on developmental rate (*p* ≥ 0.055), whereas the SDxSC line B:slow-developing parents, SDxSC line C:fast-developing parents, SDxSC line C:slow-developing parents and SCxSD line E:slow-developing parents groups displayed significant effects prior to correction for multiple tests (*p* ≤ 0.018). After correction (Bonferroni-corrected α = 4.17 × 10^-3^), only the line B:slow and line E:slow groups still showed significant effects. To investigate these results further, we repeated these analyses using only egg sacs with ≤ 40 copepodids (308 out of 360 egg sacs; ~86%) to assess if any potential density effects could be traced to large egg sacs specifically. After correction for multiple tests as above, no lines had an effect of copepod number on average egg sac developmental rate (*p* = 9.0 × 10^-3^ for SCxSD line E:slow-developing parents, and *p* ≥ 0.076 for all other groups). We also repeated our F_4_ developmental rate analysis using the egg sacs with ≤ 40 copepodids and the overall findings were unchanged (cross: *p* = 0.098; F_3_ parental developmental rate: *p* = 0.035). Therefore, taken together, our results suggested that any potential density effects on developmental rate in our study were modest, and variation in developmental rate among F_3_ *T. californicus* hybrids was inherited by their F_4_ offspring.

**Figure 6.**
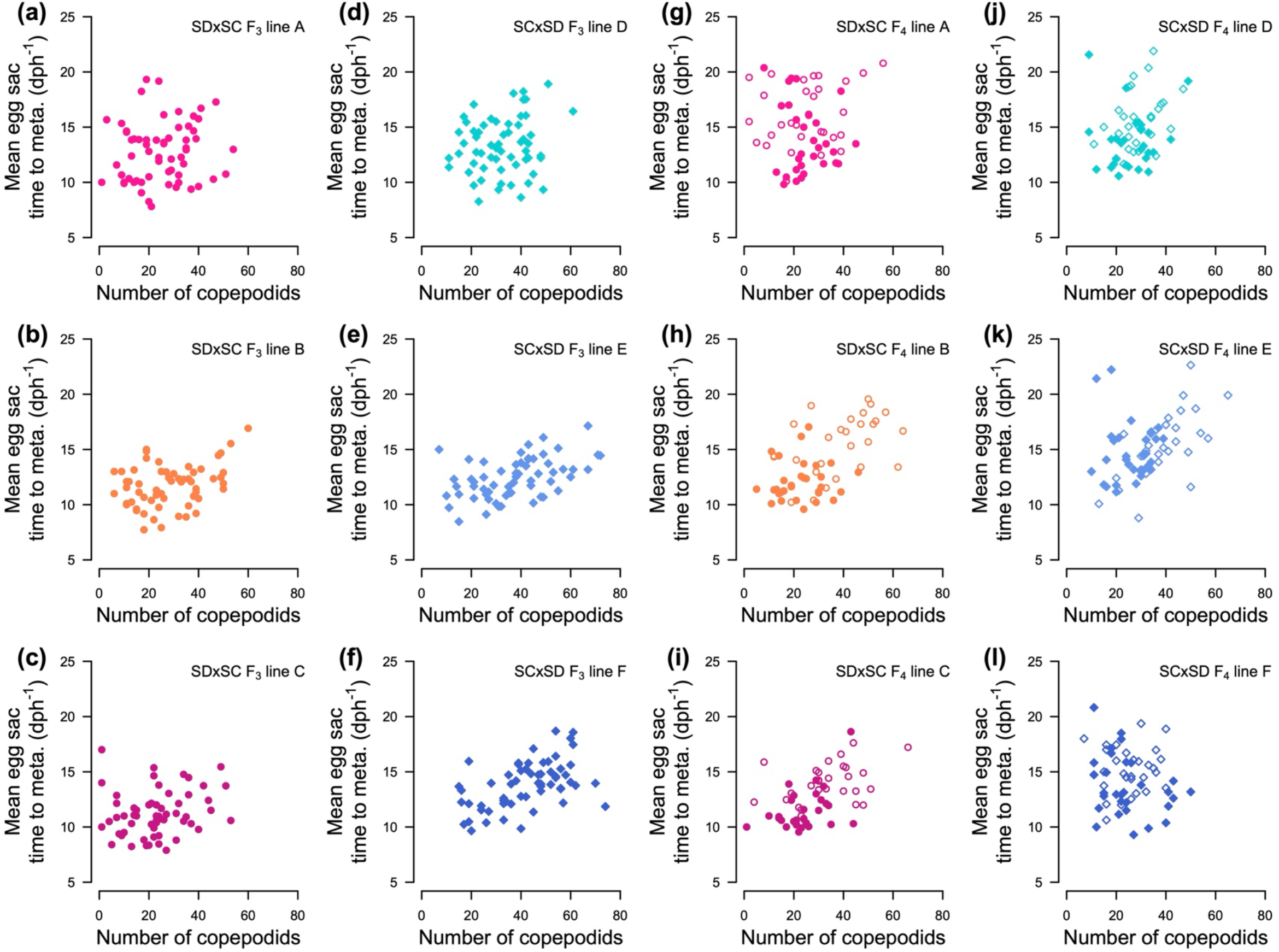
Average egg sac developmental rates plotted against the number of copepodids produced from the egg sac in F_3_ (a-f) and F_4_ (g-l, offspring of fast- or slow-developing F_3_ parents: filled or open symbols, respectively) inter-population hybrids. SDxSC crosses are shown in warm colors (a-c, g-i), and SCxSD crosses are shown in cool colors (d-f, j-l).

### Variation in ATP synthesis rates among F_4_ hybrids

Maximal *in vitro* ATP synthesis rates were measured for two sets of ETS complex I substrates: pyruvate-malate (PM), and glutamate-malate (GM). PM ATP synthesis rates in F_4_ hybrids were affected by sex (*p* < 2 × 10^-16^) and F_3_ parental developmental rate (*p* = 0.0014), but not cross (i.e., SDxSC or SCxSD; *p* = 0.82); however, there was also a significant interaction between sex and parental developmental rate (*p* = 0.040; Figure 7a). The general trends in the PM-fueled synthesis rates were similar to those for the GM ATP synthesis rates (Figure 7b), but the statistical results were somewhat different. For the GM-fueled rates, there was a significant effect of sex (*p* < 2 × 10^-16^), but not of F_3_ parental developmental rate (*p* = 0.085) or cross (*p* = 0.40), and the only significant interaction was between parental developmental rate and cross (*p* = 0.0015). Interpretation of these results is complicated by these interactions, but the clearest pattern was that of higher synthesis rates in females than in males for both the PM and GM substrates (PM: 0.0092 ± 0.0003 and 0.0049 ± 0.0001 and GM: 0.0097 ± 0.0003 and 0.0054 ± 0.0001, μ ± SEM, ATP rate:CS rate for females and males, respectively; note that data from F_4_ hybrids from the SCxSD line F:slow-developing parents group were excluded from the GM comparison here, see Methods and materials for details).

**Figure 7.**
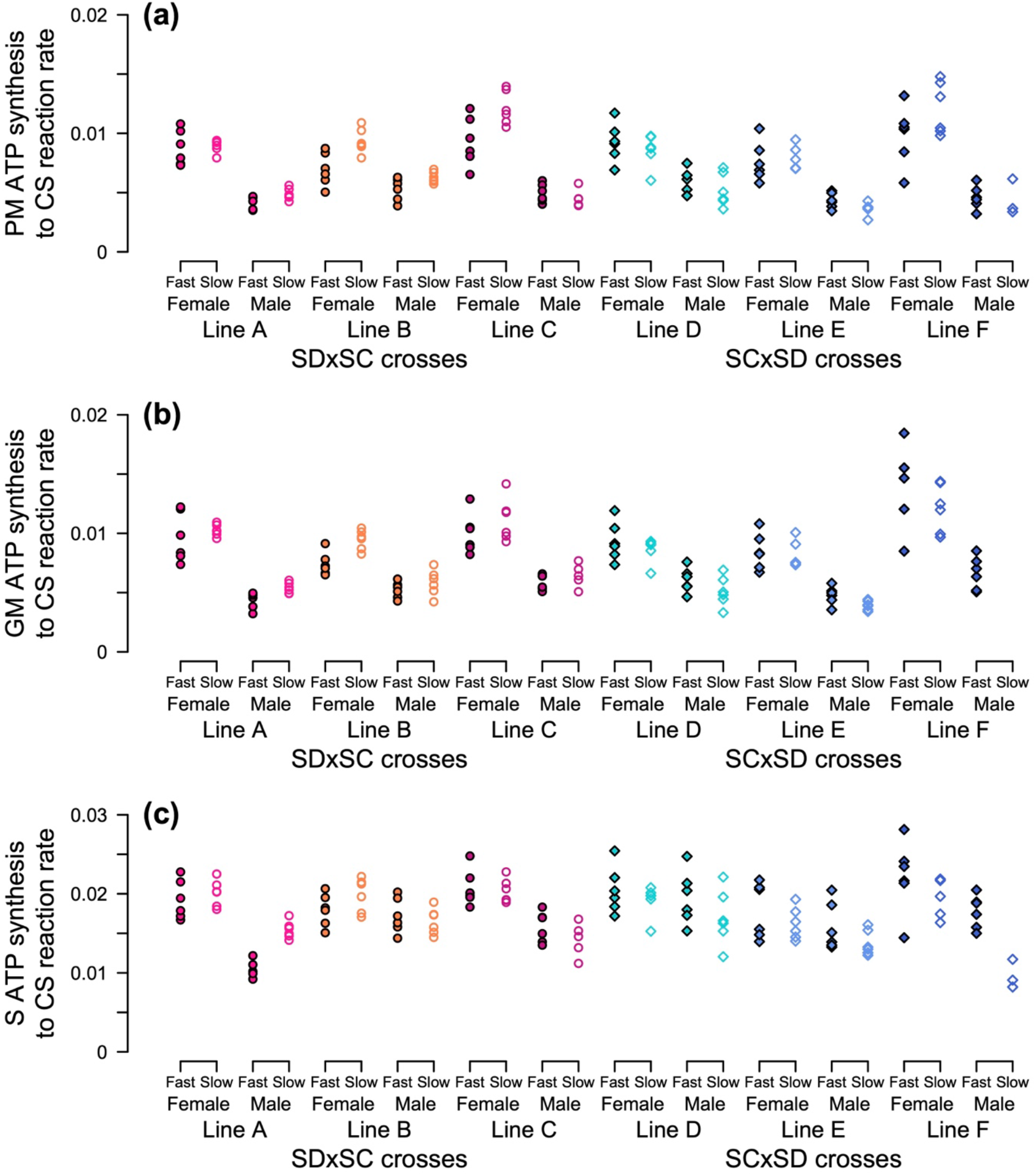
Plots of maximal *in vitro* ATP synthesis rates of mitochondria isolated from F_4_ *T. californicus* reciprocal hybrids (SDxSC, lines A-C – warm colors, circles; SCxSD, lines D-F – cool colors, diamonds) that were offspring of ‘Fast’- or ‘Slow’-developing F_3_ parents (filled or open boxes, respectively). Assays were conducted to assess electron transport system complex I-fueled (a: PM – pyruvate-malate; b: GM – glutamate-malate) and complex II-fueled (c: S – succinate) ATP synthesis under saturating conditions. Statistical results are presented in the main text of the Results.

ATP synthesis rates were also assessed using an ETS complex II substrate: succinate (S). There were significant effects of sex (*p* = 5.6 × 10^-14^) and F_3_ parental developmental rate (*p* = 1.6 × 10^-4^), but not of cross (*p* = 0.14), on the F_4_ S ATP synthesis rates (Figure 7c). Additionally, there was a significant interaction between parental developmental rate and cross (*p* = 4.6 × 10^-4^). Again, the clearest pattern was that females tended to have higher S ATP synthesis rates than males (0.0195 ± 0.0003 and 0.0156 ± 0.0004, μ ± SEM, S ATP rate:CS rate for females and males, respectively). However, the difference between females and males was proportionally smaller with the complex II substrate than with the complex I substrates, and the ratio of PM-fueled synthesis rate to S-fueled synthesis rate was higher in females than in males (sex: *p* = 5.0 × 10^-14^, F_3_ parental developmental rate: *p* = 1.3 × 10^-6^, cross: *p* = 0.75; Table 3), although as for the ATP synthesis rates independently, there were also interactive effects on the PM:S rate ratios (sex x F_3_ parental developmental rate: *p* = 0.026, sex x cross: *p* = 0.028).

**Table 3.** Ratios of pyruvate-malate (PM) ATP synthesis rate to succinate (S) ATP synthesis rate for female and male F_4_ inter-population SDxSC and SCxSD hybrids that were offspring of either fast- or slow-developing F_3_ parents.

| Cross | Line | F<sub>3</sub> parental<br>developmental<br>rate | PM:S ATP synthesis rate ratio<br>($\mu \pm \text{SEM}$) | |
| --- | --- | --- | --- | --- |
|  |  |  | Female | Male |
| SDxSC | A | *Fast* | 0.46 ± 0.02 | 0.39 ± 0.02 |
| | | *Slow* | 0.45 ± 0.01 | 0.32 ± 0.02 |
| | B | *Fast* | 0.39 ± 0.02 | 0.31 ± 0.01 |
| | | *Slow* | 0.48 ± 0.03 | 0.38 ± 0.01 |
| | C | *Fast* | 0.45 ± 0.03 | 0.31 ± 0.02 |
| | | *Slow* | 0.60 ± 0.03 | 0.32 ± 0.02 |
| SCxSD | D | *Fast* | 0.45 ± 0.01 | 0.30 ± 0.01 |
| | | *Slow* | 0.44 ± 0.02 | 0.30 ± 0.01 |
| | E | *Fast* | 0.43 ± 0.02 | 0.28 ± 0.01 |
| | | *Slow* | 0.48 ± 0.01 | 0.27 ± 0.01 |
| | F | *Fast* | 0.44 ± 0.01 | 0.26 ± 0.01 |
| | | *Slow* | 0.61 ± 0.04 | 0.45 ± 0.04 |

## Discussion

The two major findings of our study are that developmental rates do not differ between female and male *T. californicus* hybrids at any stage of development, and that phenotypic differences in F_3_ developmental rate among individuals are heritable, affecting variation in developmental rate among F_4_ hybrids. Given previous results demonstrating substantial differences in mitonuclear genotypes between fast- and slow-developing *T. californicus* hybrids (Healy & Burton, 2020; Han & Barreto, 2021), the results of the current study imply not only that these strong effects likely have similar impacts in both sexes in *T. californicus*, but also that phenotypic variation associated with these strong effects is inherited across hybrid generations. Taken together, these findings are consistent with the possibilities for mitonuclear interactions to create effective barriers for gene flow and to contribute to the early stages of reproductive isolation between populations.

### Lack of variation in developmental rate between the sexes

*T. californicus* lacks heteromorphic sex chromosomes, and has complex genetic and environmental mechanisms underlying sex determination (Alexander et al., 2015). Phenotypic differences between females and males have been well characterized for a number of traits in this species, including lifespan, viability, fertility, stress tolerances and gene expression patterns (Willett & Burton, 2001; Willett, 2010; Leong et al., 2018; Willett & Son, 2018; Foley et al., 2019; Li et al., 2019, 2020, 2022; Flanagan et al., 2021, 2022; Watson et al., 2022), and at least some of these sex-specific patterns have also been observed in inter-population hybrids. For instance, Li et al. (2022) found that females had longer lifespans than males in F1 inter-population hybrids, and Foley et al. (2013) showed that differences in nuclear allele frequencies between reciprocal F_2_ hybrids (i.e., between alternate mitochondrial genotypes) were sex-dependent, particularly for chromosome 10. However, observed sex-specific effects in hybrid *T. californicus* are not consistently aligned with the expectations of the ‘mother’s curse’ hypothesis (e.g., Watson et al., 2022), and although previous results have indicated that variation in developmental rate is generally highly dependent on mitonuclear compatibility in *T. californicus* hybrids (Healy & Burton, 2020; Han & Barreto, 2021), variation in this trait between the sexes was virtually absent at any stage of development in the current study (consistent with the less detailed results of Burton [1990]). Similarly, variation in mitonuclear effects on developmental time between females and males is generally modest or absent in *Drosophila sp.* (Jelić et al., 2015; Mossman et al., 2016a; Erić et al., 2022) despite sex-specific effects of mitochondrial genotype on many other fitness-related traits when evaluated on common nuclear genetic backgrounds (Camus et al., 2012; Kurbalija Novičić et al., 2015; Nagarajan-Radha et al., 2019, 2020; Carnegie et al., 2021). Taken together, these results and our findings for *T. californicus* suggest that effects of mitonuclear incompatibilities on developmental rate may often be inconsistent with the expectations of the ‘mother’s curse’ hypothesis.

Variation in developmental rate between females and males was assessed across two distinct phases of development in *T. californicus:* hatch to the C1 stage and the C1 stage to adult. The lack of difference between the sexes was remarkably consistent throughout both phases of development, and there was no variation in developmental rate overall between the two reciprocal crosses. The similarities in developmental rate observed here between the SDxSC and SCxSD hybrids are consistent with previous studies in *T. californicus* (Ellison & Burton, 2008b; Healy & Burton, 2020, 2022). In contrast, an unexpected result in the current study was a lack of correlation between the rates of development from hatch to C1 and from C1 to adult across the F_2_ egg sacs, which suggests that fast developers through one phase of development may not necessarily also be fast developers through the other phase. Healy and Burton (2020) found strong effects of mitonuclear compatibility on hatch to C1 developmental rate among F_2_ hybrids between the SD and SC populations; consequently, we anticipated that individuals with compatible genotypes that develop at fast rates during naupliar stages would also have fast development through copepodid stages. However, mitonuclear incompatibilities can have differential physiological effects over an organism’s life cycle (Hoekstra et al., 2018), and environmental effects may shape the consequences of mitonuclear interactions as well (Hoekstra et al., 2013, 2018; Baris et al., 2016; Mossman et al., 2016a, 2017; Drummond et al., 2019; Rand & Mossman, 2020; Rand et al., 2022). Therefore, it is possible that changes in metabolic demand throughout development or the change in holding conditions at the C1 stage in our experiment (which was necessary to confidently examine C1 to adult developmental rate) may have altered the consequences of mitonuclear incompatibilities among the individuals in our study leading to a disassociation of developmental rates before and after C1 metamorphosis. Regardless, as development from hatch to C1 and from hatch to adult have both been used to measure developmental rate in *T. californicus* (e.g., Burton, 1990; Harada et al., 2019), the lack of correlation between the two phases of development is of particular note for considering results across published studies and for designing future experiments with this species.

The role of mitonuclear interactions underlying variation in developmental rate among hybrid individuals in *T. californicus* highlights another intriguing pattern in our data, as the ATP synthesis rates for mitochondria isolated from female copepods were higher than those for mitochondria isolated from male copepods in the F_3_ to F_4_ experiment, whereas there was no difference in developmental rate between the sexes in the F_2_ experiment. Somewhat contradictory to these results, higher ATP synthesis rates are associated with fast development among F_2_ hybrid individuals in *T. californicus* (Healy & Burton, 2020; Han & Barreto, 2021). These differences may simply be a consequence of comparing results from F_2_ and F_4_ hybrids in the current study, or it is possible that metabolic demands are higher in female copepods than in male copepods during development, such that elevated ATP synthesis capacities could be a compensatory physiological response in females. However, at least in adults, there is little evidence for higher metabolic rates in female than in male *T. californicus* under control conditions in the laboratory (Powers et al., 2022).

Previous studies have detected some evidence for variation in complex IV activities between female and male *T. californicus* (Edmands & Burton, 1998, 1999), and higher ATP synthesis rates in females than in males in the current study were independent of the use of complex I or II substrates to fuel the ETS. However, the proportional increase in ATP synthesis rates from males to females was greater with complex I than complex II substrates. This is clearly evident through the elevated ratios of PM ATP synthesis rates to S ATP synthesis rates in females compared to males, and implies that oxidative phosphorylation may be more reliant on complex I function in female than in male *T. californicus*. Interacting subunits of complex I are likely sources of mitonuclear incompatibilities (e.g., Pichaud et al., 2019; Moran et al., 2021), and thus increased contributions of complex I respiration in females could also increase the susceptibility of females to mitonuclear incompatibilities. Alternatively, following from the expectations of a ‘mother’s curse’, decreased reliance on complex I respiration in males could partially negate the possibility of strong effects of mitonuclear incompatibilities on complex I function in this sex. In either case, variation in the proportional contributions of complex I and II respiration between females and males may contribute to the sex-specific effects of incompatibilities on viability, lifespan or longevity reported elsewhere (Willett & Burton, 2001; Li et al., 2022; Watson et al., 2022).

### Heritable variation in developmental rate among hybrids

There was substantial variation in developmental rate among the F_3_ hybrid individuals in our study with metamorphosis to the C1 stage occurring between 7 and 45 dph across lines. Healy & Burton (2020) observed a similar range of time to metamorphosis among F_2_ hybrids between SD and SC *T. californicus,* whereas in the current study the times to C1 metamorphosis in F_2_ hybrids were much narrower: 8 to 17 dph. The cause of this difference between the studies is unclear as culturing temperatures, salinities and photoperiods were the same, but it is possible that uncontrolled differences in culture lighting or algal growth contribute to this variation. Regardless, the similarities between the ranges of F_3_ developmental rates presented here and the ranges of F_2_ developmental rates in Healy and Burton (2020), taken together with similar average fitness consequences of mitonuclear incompatibilities in F_2_ and F_3_ *T. californicus* hybrids (Ellison & Burton, 2008b), suggest that variation in mitonuclear compatibility is likely associated with variation in developmental rate among the F_3_ individuals in the current study.

Heritability of developmental rate or growth during development has been observed in many species (e.g., *D. melanogaster* – Chippindale et al., 1997, 2004, and hybrid *Crassostrea gigas* – Pace et al., 2006; Meyer & Manahan, 2010), and given that mitonuclear effects on phenotypic variation are inherently genetic, heritable variation associated with inter-genomic interactions is expected. Consistent with this expectation, inheritance of developmental rate between F_3_ and F_4_ hybrids was observed in at least 5 of our 6 lines. Previously published studies in *T. californicus* have compared high- and low-fitness hybrid lines with variation in the extent of mitonuclear compatibility among lines (Edmands et al., 2009; Barreto & Burton, 2013; Pereira et al., 2014, 2021; Powers et al., 2021), providing indirect evidence for heritability of mitonuclear effects. However, the current study not only demonstrates the extent of this effect among hybrid individuals between successive generations (h^2^ = 0.16 ± 0.10 and 0.29 ± 0.06 for fast and slow developers, respectively), but also assesses heritability specifically for inter-individual trait variation associated with strong selection for compatible mitonuclear genotypes among hybrids (Healy & Burton, 2020; Han & Barreto, 2021).

Inheritance of F_3_ parental developmental rates in F_4_ offspring was asymmetric for fast- and slow-developing hybrids. Over all the hybrids lines, the increase in developmental rate from the average F_3_ developmental rate was 0.55 ± 0.37 d (μ ± SEM) for F_4_ offspring of fast developers compared to a decrease of −1.78 ± 0.55 d for F_4_ offspring of slow developers. Hybrid generation only had a significant effect on average developmental rate between the F_3_ hybrids and the F_4_ offspring of slow developers (see Figure 5), which is consistent with the trend for higher realized heritability in slow-developing than in fast-developing hybrids. We hypothesize that this asymmetry in response to selection may reflect the high degree of compatible mitonuclear genotypes across loci in fast-developing hybrids between the SD and SC populations; fast development requires avoiding multiple mitonuclear incompatibilities, whereas a range of possible incompatibilities may result in slow development (Healy & Burton, 2020). This suggests that there are fewer fast-developing genotypes than slow-developing genotypes, which potentially reduces additive genetic variance in fast developers and could contribute to the asymmetric response to selection between the F_3_ and F_4_ generations in the current study.

Despite the heritability of F_3_ parental developmental rates in F_4_ offspring, there was no variation in maximal ATP synthesis rates between F_4_ offspring of fast- or slow-developing parents. This lack of relationship between developmental rate and mitochondrial performance is surprising in *T. californicus*, as variation in developmental rate in inter-population hybrids has been associated with differences in mitochondrial ATP synthesis capacities both among lines (Ellison and Burton, 2006, 2008b) and among individuals (Healy & Burton, 2020; Han & Barreto, 2021). However, there was overlap in the ranges of developmental rates for the F_4_ offspring of fast- or slow-developing parents in the current study, and mitochondria were isolated from pools of copepods selected haphazardly from each F_4_ group. Therefore, it is possible that we were unable to resolve any potentially subtle differences in mitochondrial performance between the groups after the effects of allelic reassortment between the F_3_ and F_4_ generation.

## Conclusion

Genetic incompatibilities between independent taxa are key factors leading to reproductive isolation; however, the contributions of mitonuclear interactions to these effects in eukaryotes is a matter of some debate (e.g., Hill, 2017; Sloan et al., 2017; Burton, 2022). Mitonuclear incompatibilities may play disproportionately large roles in outbreeding depression and the early-stages of reproductive isolation (Burton & Barreto, 2012), particularly in eukaryotes without heterologous sex chromosomes (Lima et al., 2019), and the involvement of these incompatibilities in speciation has been proposed (Hill, 2017). These potential effects require strong selection to maintain compatible mitonuclear genotypes in hybrid eukaryotes, and our results provide evidence consistent with this selection by revealing consistent effects of incompatibilities in both sexes, and by demonstrating heritability of fitness-related trait variation (i.e., variation in developmental rate) that has previously be shown to be associated with differences in mitonuclear compatibility.

## Acknowledgements

The current study was funded by National Science Foundation grants to RSB (DEB1556466 and IOS1754347).

## Data availability

Prior to final publication, all datasets used in the current study will be submitted to the Dryad Digital Respository.

## Author contributions

TMH and RSB conceived and designed the experiments; TMH and ACH collected the data; TMH performed the analyses; TMH prepared the tables and figures, and TMH, ACH and RSB wrote the manuscript.

